# On the evolution of host specificity: a case study of helminths

**DOI:** 10.1101/2021.02.13.431093

**Authors:** Alaina C. Pfenning-Butterworth, Sebastian Botero-Cañola, Clayton E. Cressler

## Abstract

The significant variation in host specificity exhibited by parasites has been separately linked to evolutionary history and ecological factors in specific host-parasite associations. Yet, whether there are any general patterns in the factors that shape host specificity across parasites more broadly is unknown. Here we constructed a molecular phylogeny for 249 helminth species infecting free-range mammals and find that the influence of ecological factors and evolutionary history varies across different measures of host specificity. Whereas the phylogenetic range of hosts a parasite can infect shows a strong signal of evolutionary constraint, the number of hosts a parasite infects does not. Our results shed new light on the evolution of host specificity in parasites, suggesting that phylogenetic breadth may capture the evolutionary potential of a parasite to jump between hosts, whereas the number of hosts may reflect ecological opportunity. Finally, we show parasite phylogenies can also provide an alternative perspective on zoonosis by identifying which hosts are infected by a broad phylogenetic range of parasites.

## INTRODUCTION

Parasites vary considerably in their host specificity, or the range of hosts they can infect, and previous work has identified a large number of ecological and evolutionary factors that may lead to variation in host specificity (Bernays and Graham 1988; Poulin 1992; Desdevises et al. 2002; Mouillot et al. 2006; Clark and Clegg 2017). In some systems, host specificity appears to be more strongly shaped by local environment (e.g., host diversity/abundance and abiotic factors, Krasnov et al. 2005; Loiseau et al. 2012; Dallas and Presley 2014), whereas in other systems, host specificity appears to be constrained by evolutionary history (e.g., outcomes of adaptive evolution, Desdevises et al. 2002; Clark and Clegg 2017). These contrasting results are perhaps not surprising, given the phylogenetic, phenotypic, and ecological diversity of parasites – perhaps there are no general patterns that underlie the evolution of specificity across parasite lineages. However, there is value in exploring that question, as a broader understanding of the ecological and evolutionary factors that shape host specificity may help to identify hosts and parasites that are the most likely reservoirs of novel zoonoses.

Studies like those referenced above have assessed the determinants of variation in host specificity using taxonomically restricted datasets that focus on a single group of parasites (e.g., fleas, avian malaria, etc.). This taxonomic restriction can allow researchers to test specific hypotheses (e.g., whether host specificity varies with geography; Krasnov et al. 2005; Krasnov et al. 2008; reviewed in Poulin et al. 2011) and to more readily identify ecological and physiological factors likely to be driving differences in specificity among parasite species (e.g., whether host specificity is shaped by host phylogeny; McCoy et al. 2001; Fallon et al. 2005; Clark and Clegg 2017). However, the widespread variation in host specificity among groups of parasites suggests that inferences that apply to one group will likely be of limited value in other groups. For instance, monogeneans, ecto-parasitic flat worms, have high host-specificity (~74% infect a single host, Bikhovski 1957) and are thought to have diversified during the Cretacious period (~100 mya, Kearn 1994). Monogeneans diverged from Platyhelminthes (which emerged ~270 mya, Dentzien-Dias et al. 2013), and the oldest known helminth lineages emerged ~550 mya (Zhang et al 2020). Given such ancient divergences, it would be unwise to conclude that all helminths are highly host-specific, or that the factors that drive variation in specificity among monogeneans will also be important to Platyhelminthes. Nevertheless, helminths like these are perhaps the best group of parasites to study to determine whether local environment or evolutionary history are the primary determinants of host specificity across the broadest phylogenetic scale studied to date.

Helminths are perhaps the most successful parasites in the world, in terms of global prevalence (Dallas et al. 2018). They are a diverse group consisting of four major groups of parasitic worms—acanthocephalans, cestodes, nematodes, and trematodes—that differ in their morphology, transmission, and host specificity (Mackiewicz 1988; Hayunga 1991; Kennedy 2006). They also differ in their evolutionary history (Weinstein and Kuris 2016); for example, nematodes coevolved alongside vertebrates (Poinar 2011), whereas cestodes have emerged more recently (Baer 1952). Previous studies indicate that variation in life history among the helminth groups may help explain key differences in host specificity (Stunkard 1957; Pedersen et al. 2005). For instance, the generalism observed in acanthocephalans is thought to be due to the presence of a free-living stage in their life cycle (Stunkard 1957). The phylogenetic and ecological diversity of helminths presents a rare opportunity to discern whether there are fundamental processes that determine host specificity across broad taxonomic and phylogenetic scales.

Here, we construct a molecular helminth phylogeny for 249 species of helminths that parasitize free-living mammals. This novel phylogeny is the largest ever built for this group, and thus provides unprecedented insight into the factors that shape the evolution of host specificity. We use this phylogeny to test the evidence for two hypotheses in shaping variation in host specificity: (1) evolutionary history, where closely related helminths would always have similar host specificities, and (2) local ecology, where closely related helminths living in different environments would have very different host specificities. We tested these hypotheses using two metrics of host specificity: the number of hosts infected (taxonomic breadth) and the mean pairwise phylogenetic distance among hosts (MPD, Webb et al 2002). We find that the mean pairwise phylogenetic distance among hosts is shaped by evolutionary history, whereas taxonomic breadth is not. Using this phylogeny, we also assessed which mammal species may be more likely to serve as reservoirs for emerging infectious diseases finding that Old and New World monkeys host a phylogenetically diverse set of helminths making them potential sources of zoonotic disease.

## MATERIAL AND METHODS

### Host-parasite databases

We obtained records of mammal-helminth interactions from two datasets: Global Mammal Parasite Database (GMPD, Stephens et al. 2017) and a novel dataset assembled from parasitology museums (Botero-Cañola 2020). The GMPD includes taxonomic and trait data obtained from the literature for over 10,000 host-helminth associations of wild populations from four Orders of mammals: carnivores (Carnivora), primates (Primates), and ungulates (Artiodactyla and Perissodactyla). Some limitations of the GMPD are the exclusion of rodent hosts and the lack of expert identification of helminths, which could lead to an overestimation of the number of specialist helminths due to missing hosts or the misidentification of rare species. To account for these potential sources of bias we also ran all analyses with the dataset assembled from voucher specimen-verified museum collections. This dataset includes host-parasite associations for Nearctic mammals in the Orders Artiodactyla (even-toed ungulates), Carnivora (carnivores), Didelphimorphia (opposums), Eulipotyphla (moles and shrews), Lagomorpha (rabbits), and Rodentia (rodents) obtained from the H.W. Manter Parasitology Collection, the former United States National Parasitology Collection, the Museum of Southwestern Biology, and the Canadian Museum of Nature.

### Phylogenetic construction

We searched Genbank for DNA sequences (CO1, 18S, and 28S) of all 920 species of helminths included in the GMPD (Stephens et al. 2017), resulting in 249 species. We combined slow evolving sequences (18S and 28S) that resolve deep phylogenetic relationships with a fast-evolving sequence (CO1) to resolve recent relationships. Sequences were aligned using Multiple Sequence Alignment (MUSCLE v. 3.8.425, Edgar 2004) and manually edited in Geneious Prime 2019.0.4 (Kearse et al. 2012). The sequences for each species included 8848 base pairs—1707, 3550, 3591 base pairs for CO1, 18S, and 28S respectively. We implemented a Bayesian likelihood analysis on the combined data set with a GTR + G + I substitution model (PartitionFinder v.2.1.1; Lanfear et al. 2016) and a relaxed lognormal molecular clock model selected in BEASTv.2.5.2 (Bouckaert et al. 2014). We ran three replicates of the Bayesian Markov chain Monte Carlo (MCMC) analysis for 60 million generations each, sampling every 1000 generations (BEASTv.2.5.2; Bouckaert et al. 2014). We assessed the MCMC log files for convergence (effective sample size, ESS > 200, Tracer v.1.7.1; Rambaut et al. 2018) and removed the first 44% as burn-in (TreeAnnotator v.2.5.2, Bouckaert et al. 2014). The constructed tree is consistent with the topology of published trees for smaller helminth groups (Acanthocephala—Vereweyen et al. 2011; García-Varela et al. 2013, Nematoda—Blaxter et al. 1998; Nadler and Hudspeth 2000, Platyhelminthes—Knapp et al. 2015; Sharma et al. 2016) and clearly assigns recognized Orders to monophyletic clades (Supplemental Figure 1). Focusing only on the host-parasite associations for which we have phylogenetic information for the parasite, the analyses of the GMPD data included 971 associations among 197 mammal species and 249 helminth species; the analyses of the Nearctic museum data included 363 associations among 67 mammal species and 90 helminth species.

### Phylogenetic signal in host specificity metrics

Metrics of host specificity used here—taxonomic breadth and mean pairwise phylogenetic distances (MPD)—were calculated following Park et al. (2018). Taxonomic breadth refers to the number of hosts a parasite can infect (Rhode 1980); whereas mean pairwise phylogenetic distance refers to the mean branch length of all pairs of host species infected by a given parasite (Harnos et al. 2017). We used the Phylogenetic Atlas of Mammal Macroecology’s mammal phylogeny (PHYLACINE; Faurby and Svenning 2015; Faurby et al 2018) to determine the MPD of hosts for each helminth using the R package *picante* (Kembel et al. 2010). For single host parasites, we set host taxonomic breadth equal to one and MPD equal to 0. However, these parasites may have a disproportionate influence on our phylogenetic conclusions: for example, moving from one host to two can have a dramatic effect on MPD, in particular, and high host specificity may be a result of under sampling rather than true specificity. To determine how sensitive the analyses were to the inclusion of single host parasites we reran all analyses, excluding single host parasites.

We assessed whether MPD and taxonomic breadth exhibited phylogenetic signal by testing whether a non-phylogenetic model (white noise) or Pagel’s lambda best explained the data (Pagel 1999). The non-phylogenetic model (white noise) treats each species MPD or taxonomic breadth as an independent sample drawn from null distribution with shared mean and variance. High support for the non-phylogenetic model would indicate that the trait is not constrained by phylogenetic relationships. Under Pagel’s lambda, the strength of MPD or taxonomic breadth’s phylogenetic signal is estimated between 0 and 1 (where 0 indicates that the measure has been evolving as if the species were related by a “star” phylogeny, and 1 indicates that the measure has been evolving under Brownian motion, BM; Felsenstein 1973). High support for a Pagel’s lambda model greater than 0 would indicate that closely related helminths have more similar MPD or taxonomic breadth than distantly related helminths.

### Evolutionary model fitting

Based on the results of the phylogenetic signal analyses, we used phylogenetic comparative analysis to further explore the evolutionary history of MPD, with and without single host parasites. Specifically, we fit the MPD data to the parasite phylogeny under different models of trait evolution: Brownian motion (BM), a single regime Ornstein-Uhlenbeck (OU1), and a multiple regime OU based on Phyla. The BM model assumes neutral trait evolution, with a variance between taxa that is proportional to the phylogenetic distance (i.e., the branch lengths) between the taxa (Felsenstein 1973; Felsenstein 1985). In contrast, the OU1 model incorporates a deterministic trend towards a single trait value, often interpreted as stabilizing selection towards an optimum (Hansen 1997; Butler and King 2004). The OU model can also account for multiple selective regimes, allowing for different trait optima in each regime (Hansen 1997; Butler and King 2004; Beaulieu et al 2012); this is often used to test whether ecological or environmental differences among species influence trait evolution after accounting for phylogeny (e.g., the influence of trophic transmission on the life histories of helminth parasites: Benesh et al. 2011, 2014). We tested an OU model with three regimes corresponding to the different parasite Phyla. All models were fit using the R package ouch, with fit assessed using *ω*AICc (R v.3.5.2, R Core Team 2018; Burnham and Anderson 2004; Butler and King 2004; Cressler et al. 2015).

To assess the support for the best-fitting evolutionary model, we simulated 1000 datasets with the best-fitting parameter estimates for each model and then re-fit the competing models to the simulated data to create distributions of likelihood values under different generating models (Boettiger et al. 2012). We used a Phylogenetic Monte Carlo (Boettiger et al. 2012) approach to calculate distributions of the test statistic

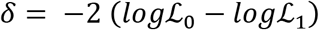

where 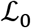 is the likelihood of the simpler model and 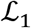 is the likelihood of the more complex model. We computed this test statistic for the BM:OU3 comparison and the OU1:OU3 comparison for the GMPD and Nearctic datasets with and without specialist parasites.

To determine whether the observed value of 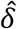 from fitting the real data was significantly different from a null expectation, we approximated a p-value (the probability of observing *δ* if the simpler model were true). We simulated 1000 datasets using the simpler model at its MLE parameter estimates and then we fit both the simple model and OU3 to the simulated dataset and computed the values of *δ*. This produces a null distribution of *δ* under the simpler model. We calculate an approximate p-value as the fraction of *δ* values in this distribution that are as extreme as the observed value, 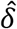.

To determine the power of each comparison, we computed the probability of rejecting the simpler model when the data were generated by the more complex (OU3) model. We simulated 1000 datasets using the OU3 model at its MLE parameter estimates and then we fit both models to the simulated dataset and computed the values of *δ*. The fraction of δ values that are larger than the 95^th^ percentile of the null distribution generated under the simpler model above gives an estimate of power.

### Phylogenetic Generalized Least Squares

We assessed whether variation host specificity (MPD) is shaped by parasite and host life history traits. Specifically, we used phylogenetic generalized least squares (PGLS) to test for if parasite length (Benesh et al. 2017), transmission mode (close-contact, environmental, and trophic transmission; Stephens et al. 2017), and average host mass (in grams, PHYLACINE, Faurby and Svenning 2015; Faurby et al 2018) explain the variation in MPD for both datasets with and without single host helminths. PGLS analyses were conducted with 52 species for which parasite body length was well documented using the R package *caper* (Orme et al. 2013). To account for phylogenetic signal, we fit each dataset’s PGLS using Pagel’s *λ* (Pagel 1999).

### Helminth phylogenetic diversity within mammal clades

We calculated the mean pairwise phylogenetic distances (MPD) of helminths that infect each host to identify patterns of helminth diversity within host clades. The helminth phylogeny constructed in this study was used to determine the MPD of helminths that infect each host, using the R package *picante* (Kembel et al. 2010). Host MPD was visualized on the mammal phylogeny using the R package *phytools* (Revell 2012). We also determined the pairwise distance (PD) of each mammal host to humans to assess which hosts are closely related to humans and are also infected by a broad range of helminths, potentially identifying host species at high risk for emerging infectious disease.

## RESULTS

### Phylogenetic signal in host specificity metrics

We assessed whether the host specificity metrics, MPD and taxonomic breadth, exhibited phylogenetic signal and found that MPD had phylogenetic signal and taxonomic breadth did not, regardless of the dataset or the inclusion of single host parasites. For MPD, datasets that excluded single host parasite had higher *λ* values, indicating stronger phylogenetic signal (Nearctic with single host parasites: *λ* = 0.38; Nearctic without single host parasites: *λ* = 0.92; GMPD with single host parasites: *λ* = 0.61; GMPD without single host parasites: *λ* = 0.90). For taxonomic breadth, the *λ* value did not change with the exclusion of single host parasites (Nearctic with single host parasites: *λ* = 0.09; Nearctic without single host parasites: *λ* = 0.09; GMPD with single host parasites: *λ* = 0.00; GMPD without single host parasites: *λ* = 0.00).

Given that MPD showed phylogenetic signal, MPD values were visualized on the helminth phylogeny, including single host helminths, for the GMPD and Nearctic dataset (Figures 1 and 2, respectively). Within the GMPD phylogeny, there are clades of specialist helminths (e.g., *Mansonella sp.* and *Cylicocyclus sp.,* Figure 1) and clades of generalist helminths (e.g., *Alaria sp.* and *Trichinella sp.,* Fig. 1). There are also clades in which one or a few species’ host specificity varies from the majority of the clade (e.g., *Taenia madoquae* is a specialist in a predominantly generalist clade, Figure 1). Overall, there is no obvious sign of evolution either towards or away from specialism across the tree. The Nearctic phylogeny show similar variation, including clades of only specialist helminths (e.g., *Trichuris sp.,* Figure 2) and clades with both extreme specialists and generalists (e.g., acanthocephalans, Figure 2).

**Figure 1.**
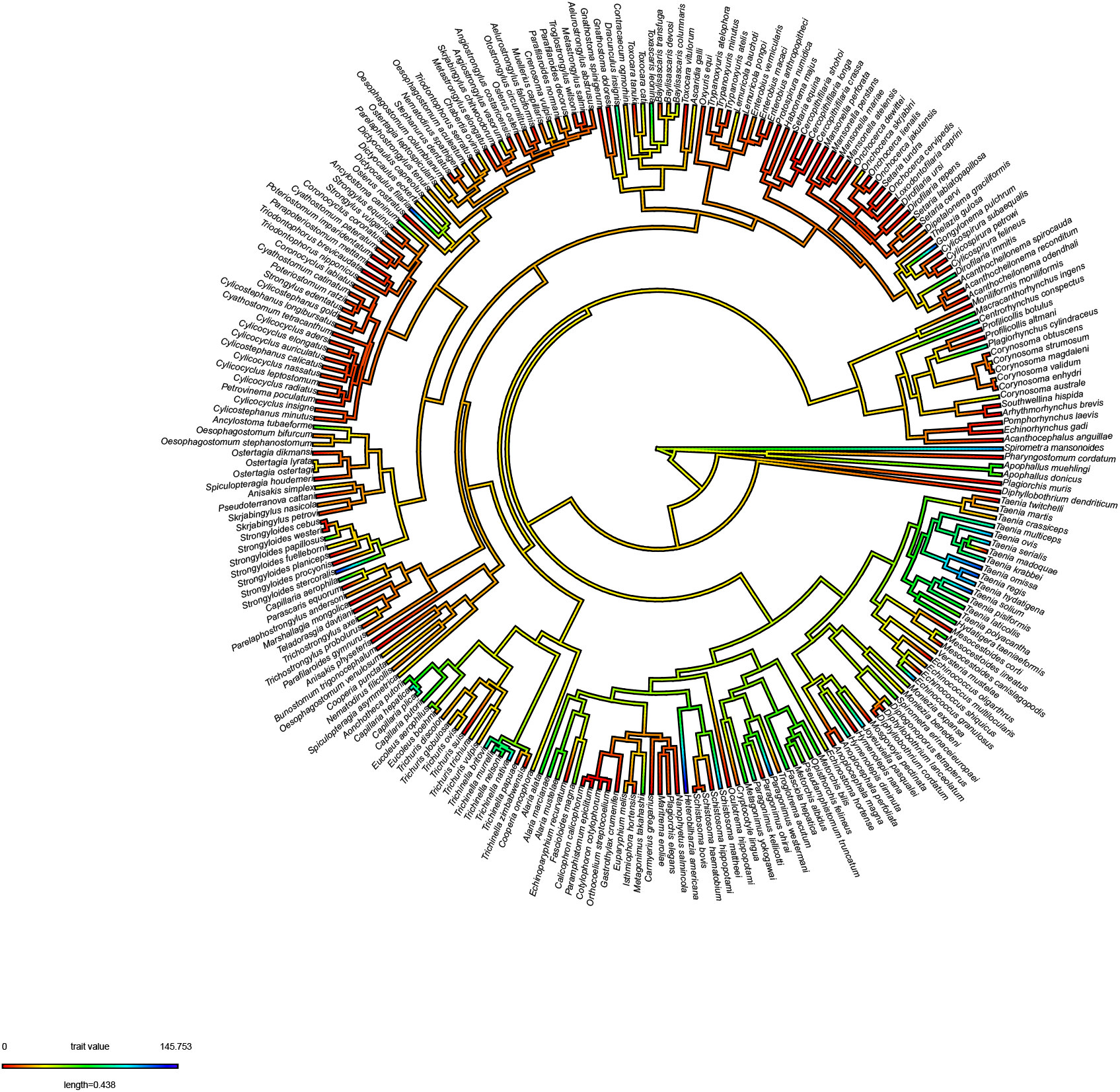
GMPD helminth phylogeny colored by MPD (warmer colors indicate higher specialism). Single host helminths are included.

**Figure 2.**
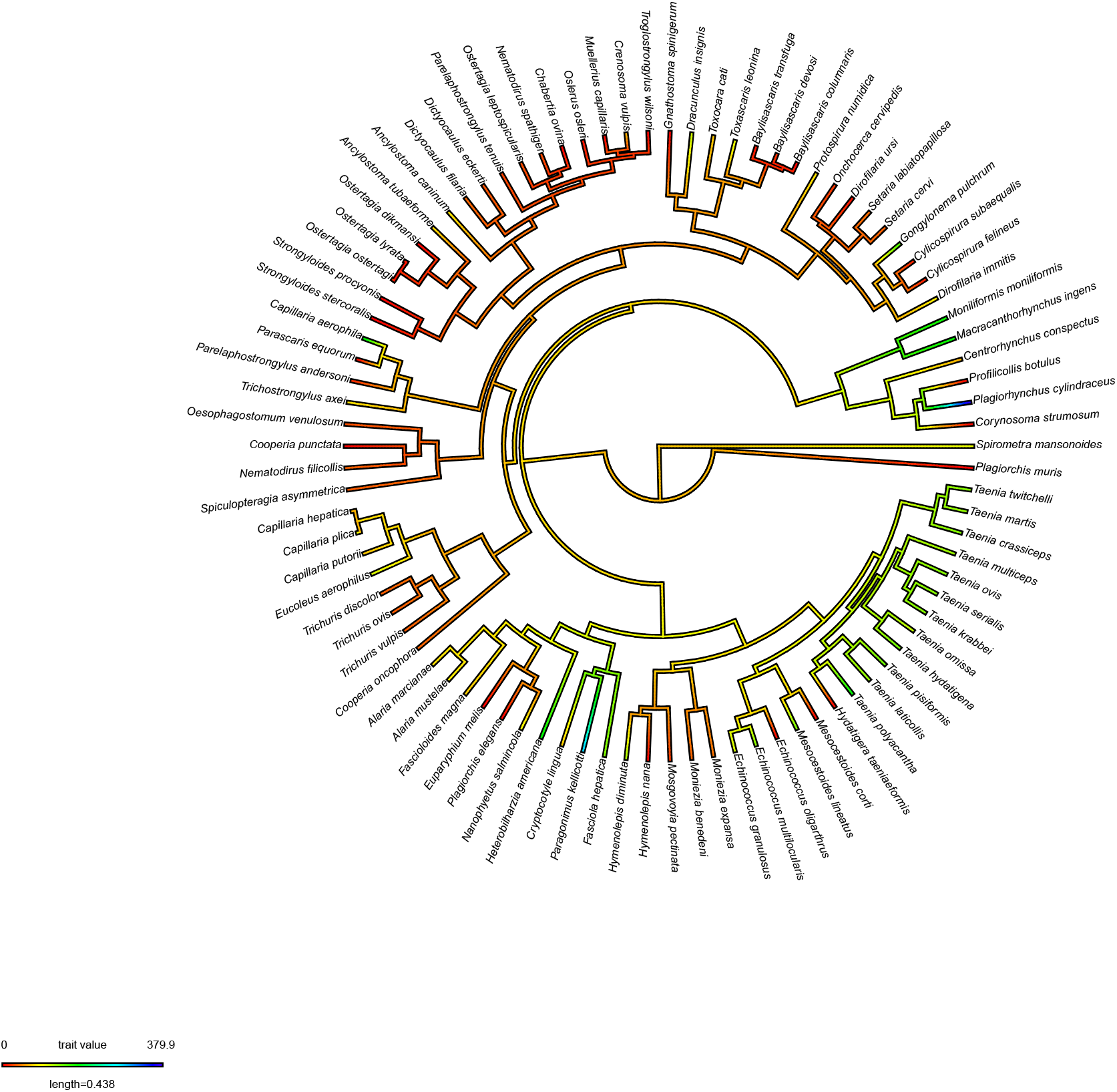
Nearctic helminth phylogeny colored by MPD (single host helminths included).

### Evolutionary model fitting

We modelled the evolutionary history of host specificity (MPD) and established that an OU3 model with regimes for each of the parasite Phyla best described the Nearctic and GMPD dataset, with and without the exclusion of single host parasites (using *ϖ*AICc; Table 1; full details in Supplemental Tables 1-4). We assessed the support for the OU3 model using a Phylogenetic Monte Carlo approach. There was high support for the OU3 model compared to the BM model for both datasets, with and without single host parasites (Supplemental Figure 2). The OU3 model had high support compared to the OU1 model for both datasets, with single host parasites. However, the GMPD dataset without single host parasites had lower power, but still maintained a high p-value, suggesting that it was unlikely that the data was generated by an OU1 model (Supplemental Figure 3).

**Table 1.** Weighted AICc scores for each of the three evolutionary models (a three-regime OU model [OU3], a single-regime OU model [OU1], and a Brownian motion model [BM]) fit to the MPD data for each dataset, with and without specialist helminths.

| **Model** | **Nearctic** | **Nearctic**<br>**(no specialists)** | **GMPD** | **GMPD**<br>**(no specialists)** |
| --- | --- | --- | --- | --- |
| OU3 | 0.99 | 0.99 | 0.99 | 0.88 |
| OU | $9.9 \times 10^{-3}$ | $4.1 \times 10^{-4}$ | $6.5 \times 10^{-3}$ | 0.11 |
| BM | $2.2 \times 10^{-7}$ | $5.3 \times 10^{-5}$ | $1.6 \times 10^{-20}$ | $5.9 \times 10^{-6}$ |

The OU3 model parameter estimates for the GMPD and Nearctic datasets, with and without single host parasites, can be found in Table 2. For both datasets, with and without single host parasites, the nematode optimum is at a lower MPD value, indicating higher host specificity for nematodes than Platyhelminthes and Acanthocephala. In the GMPD, with and without single host parasites, the Platyhelminthes optimum is the largest, whereas in the Nearctic dataset, the Acanthocephala optimum is the largest. However, regardless of dataset, the stationary variance of the OU process (given by *σ*^2^/2*α*) is very large, indicating that, within each taxonomic clade, there is considerable variation in MPD among parasite species.

**Table 2.**
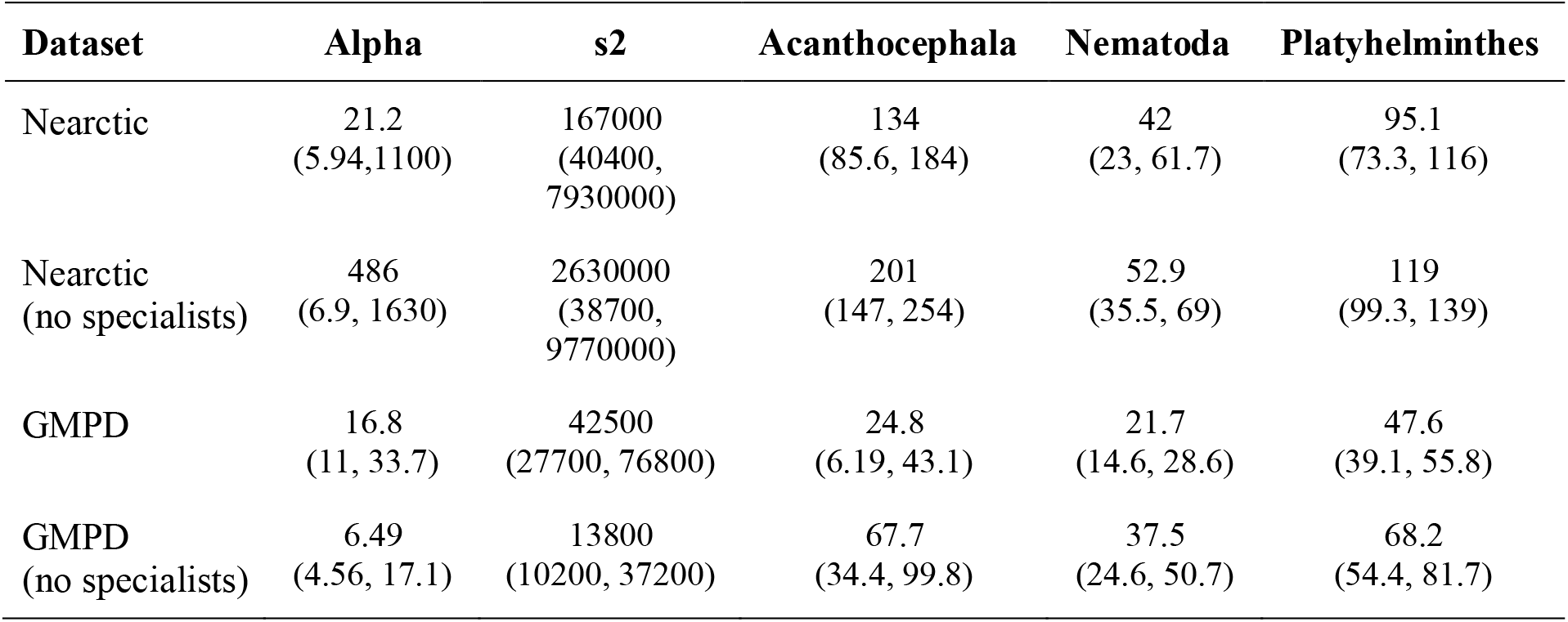
Parameter estimates (confidence intervals) for three-regime OU models (OU3) fit to the MPD data for each dataset, with and without specialist helminths.

| Dataset | Alpha | s2 | Acanthocephala | Nematoda | Platyhelminthes |
| --- | --- | --- | --- | --- | --- |
| Nearctic | 21.2<br>(5.94, 1100) | 167000<br>(40400, 7930000) | 134<br>(85.6, 184) | 42<br>(23, 61.7) | 95.1<br>(73.3, 116) |
| Nearctic<br>(no specialists) | 486<br>(6.9, 1630) | 2630000<br>(38700, 9770000) | 201<br>(147, 254) | 52.9<br>(35.5, 69) | 119<br>(99.3, 139) |
| GMPD | 16.8<br>(11, 33.7) | 42500<br>(27700, 76800) | 24.8<br>(6.19, 43.1) | 21.7<br>(14.6, 28.6) | 47.6<br>(39.1, 55.8) |
| GMPD<br>(no specialists) | 6.49<br>(4.56, 17.1) | 13800<br>(10200, 37200) | 67.7<br>(34.4, 99.8) | 37.5<br>(24.6, 50.7) | 68.2<br>(54.4, 81.7) |

### Life-history traits and host specificity

For each dataset, with and without specialist helminths, we assessed whether variation host specificity (MPD) was shaped by parasite and host life history traits (parasite length, close-contact transmission, environmental transmission, trophic transmission, and average host mass) using PGLS. In the Nearctic dataset, with specialists, the best model only included host mass as a predictor (R^2^ = 0.18, *p* = 0.01). We found a significant negative effect of host mass indicating that more specialist helminths are found in large-bodied mammals *(p* = 0.01, See Supplemental File). In the Nearctic dataset, without specialists, the best model included parasite length, close transmission, environmental transmission, and trophic transmission (R^2^ = 0.07, *p* = 0.27); however, none of these predictors were significant. In the GMPD, with and without specialists, the best model only included close transmission (R^2^ = 0.01, *p* = 0.24; R^2^ = 0.05, *p* = 0.09). In neither model did close transmission have a significant effect on MPD.

### Helminth phylogenetic diversity within mammal clades

For each dataset, we calculated the mean pairwise phylogenetic distance of helminths infecting each mammal and assessed patterns of helminth diversity within mammal clades. In the Nearctic dataset, 28% of the mammals were only infected by one helminth (MPD = 0), and thirteen species (19%) were infected by multiple phylogenetically distant helminths (MPD value > 0.9; since the helminth tree is scaled to a height of one, a host with parasite MPD > 0.9 indicates that its parasites diverged deep in the tree). Of mammals with high parasite MPDs, 15% are ungulates, 54% are carnivores, and 31% are rodents (Supplemental Figure 4). In the GMPD, 38% of the mammals were only infected by one helminth (MPD = 0), and 27 species (14%) were infected by multiple phylogenetically distant helminths (MPD value > 0.9). Of these, 22% are ungulates, 63% are carnivores, and 11% are primates (Supplemental Figure 5). There are five carnivores with high MPD (> 0.9) values in both datasets *(Lynx rufus, Urocyon cinereoargenteus*, *Lontra canadensis*, *Procyon lotor*, and *Ursus americanus*). *Urocyon cinereoargenteus* (gray fox) and *Procyon lotor* (raccoon) have MPD values greater than 0.95 in both datasets.

We plotted the pairwise distance (PD) of each mammal host to humans against the MPD of helminths infecting each mammal to identify mammals that might pose a higher risk for a spillover event. In the GMPD dataset, which does not include rodents, we found that new world monkeys, which are closely related to humans, have the highest MPD values (Figure 3). In the Nearctic dataset, the species with the highest MPD values (i.e., *Procyon lotor* and *Peromyscus gossypinus*) are not closely related to humans (Figure 3).

**Figure 3.**
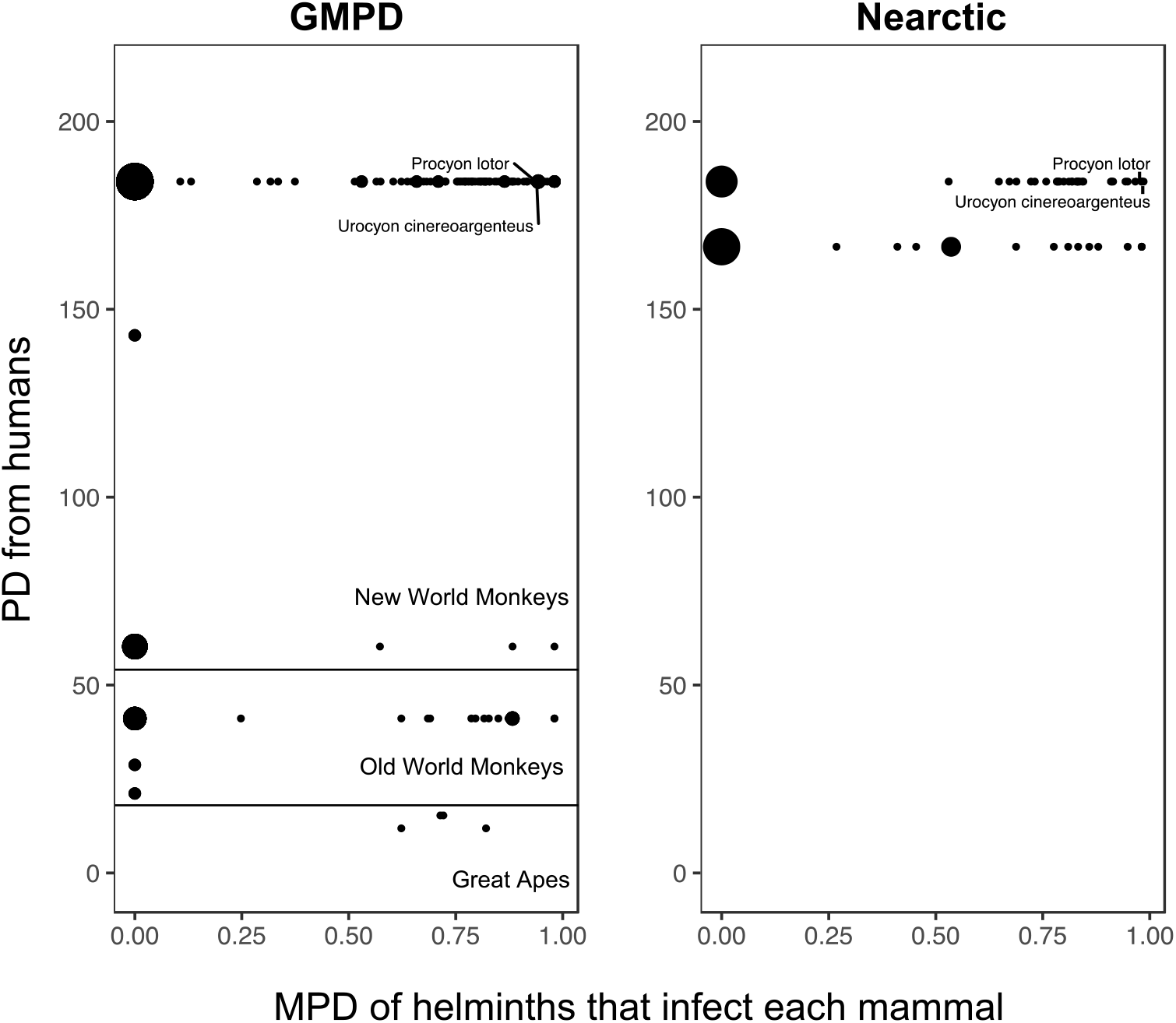
Mean pairwise phylogenetic distance (MPD) of helminths infecting each mammal compared to their phylogenetic distance to humans in the GMPD and Nearctic dataset. *Didelphis virginiana* and *Didelphis marsupialis* are most distantly related to humans (PD = 380), thus were excluded from the figure.

## DISCUSSION

Prior work has assessed the variation in host specificity in taxonomically restricted groups of parasites (Desdevises et al. 2002; Krasnov et al. 2005, 2008; Hellgren et al. 2009; Loiseau et al. 2012; Dallas and Presley 2014; Clark and Clegg 2017). These and other studies on host specificity are necessary to address specific hypotheses about the determinants of host specificity within a parasite group (e.g., determining whether monogenean host specificity varies with host body size; Desdevises et al. 2002). Krasnov et al. have studied the effects of specific environmental factors on the host specificity of fleas that infect mammals finding that fleas with a larger geographic range also have a larger host breadth and that these increase with latitude (2005; 2008). Additionally, work on two species of fleas, *Listropsylla agrippinae* and *Chiastopsylla rossi,* has indicated that host breadth varies with time spent on the host, suggesting that life history traits shape host specificity (Van der Mescht et al. 2015). In this study, we asked whether we could identify any general patterns of variation in host specificity. The general patterns indicated in this study would benefit from additional work within specific clades to identify the specific evolutionary or ecological interactions shaping host specificity.

We used a novel helminth phylogeny and two helminth-mammal databases to test competing hypotheses about the determinants of variation in host specificity. The first hypothesis we considered suggests that host specificity is determined by local environmental factors, such as host diversity and abundance or abiotic factors. Under this hypothesis, closely related parasites living in different environments would have very different host specificities, whether measured using the phylogenetic distance between hosts (MPD) or the number of hosts (taxonomic breadth). The second hypothesis suggests that host specificity is determined by evolutionary constraint. Under this hypothesis, closely related parasites would always have similar host specificities. Finally, host specificity could be determined by some combination or neither of these hypotheses.

We found, regardless of the dataset used or the inclusion of single host parasites, that MPD is constrained by evolutionary history and taxonomic breadth is not. This suggests that the phylogenetic breadth of hosts a parasite can infect is shaped by evolution, and possibly coevolution (Hadfield et al. 2014), whereas the number of hosts a parasite can infect is not. Similar results have been found in other helminths (monogeneans, Desdevises et al. 2002; trematodes, cestodes, and nematodes; Mouillet et al. 2006). This difference between host specificity metrics suggests that host specificity is determined by local environment *and* evolutionary history. One possible interpretation of these results is that the phylogenetic range of hosts a parasite can infect is constrained its life history traits (given the results of PGLS, it appears that the life history traits that effect MPD likely vary with parasite species), whereas the number of hosts that a parasite can infect within that range is determined by the availability of those hosts within the extent of the parasite’s geographic range. Some studies have shown support for this idea, finding that the host range of a parasite varies across different areas of their geographic distribution (Thompson 2005; Ricklefs 2010; Huang et al. 2018). Additional evidence for this interpretation could come from studies counting how many hosts within a parasite’s phylogenetic range have an overlapping geographic range with the parasite: this number should correspond to the parasite’s taxonomic breadth. Together, these results suggest that, like free-living species, parasites have a fundamental (shaped by MPD) and a realized (taxonomic breadth) niche.

MPD was best described by an OU3 model with regimes specified by Phyla for both datasets, with and without single host helminths, indicating that host specificity has evolved differently within clades of helminths. The OU3 model parameters for each dataset indicate that Nematodes are the most host specific phylum of helminths. However, the most generalist phylum of helminths varies with the dataset used; in the GMPD Platyhelminthes are the most generalist and in the Nearctic dataset the Acanthocephala are the most generalist. The Acanthocephala are trophically transmitted, with both a free-living life stage and life stages inside their intermediate and definitive hosts (Kennedy, 2006); given this variability in habitats and hosts, it makes some sense that they would be generalist, able to infect a broad range of hosts (Pedersen et al. 2005). Given that both Platyhelminthes and Nematodes contain a diverse range of life cycles, habitats, and hosts, further research is needed to understand why Platyhelminthes appear more generalist and Nematodes appear more specialist.

Mammals are considered common reservoirs for zoonotic pathogens because they are closely related to and have frequent interactions with humans (Brook and Dobson 2015; Han et al. 2015). Prior research has used mammal traits and parasite host specificity to predict which mammal-parasite interactions are most likely to lead to spill-over events (Stephens et al. 2016; Olival et al. 2017). These and other research indicate that parasites with low host specificity, i.e., generalists, are a greater zoonotic threat because they are more likely to infect novel hosts than specialists (Woolhouse and Gowtage-Sequeria 2005; Johnson et al. 2015).

Here we used the helminth phylogeny to add a new perspective to this discussion, by considering the phylogenetic diversity of parasites infecting the host, in addition to the host range of the parasites. Hosts that are closely related to humans and are infected by phylogenetically distance parasites may be more likely to harbor zoonotic pathogens. We found that both Old and New World monkeys serve as hosts to many distantly related helminths (Figure 3). This is consistent with prior work indicating that zoonotic diseases ‘jump’ from hosts that are closely related to humans (Pederson and Davies 2009; Han et al. 2016). This helminth phylogeny provides a unique approach for addressing which hosts are likely to harbor zoonotic diseases by allowing for the identification of host clades that are infected by phylogenetically distant parasites.

## Acknowledgements

We thank the members of the Macroecology of Infectious Disease Research Coordination Network (NSF-DEB 131223), especially J. Davies, M. Farrell, J. Herrera, A. Park, and P. Stephens, for helpful discussion and development of methods, and S. Huang for their comments on the manuscript.

